# Identification of the critical replication targets of CDK reveals direct regulation of replication initiation factors by the embryo polarity machinery in *C. elegans*

**DOI:** 10.1101/2020.06.24.168708

**Authors:** Vincent Gaggioli, Manuela R. Kieninger, Anna Klucnika, Richard Butler, Philip Zegerman

**Affiliations:** Wellcome Trust/Cancer Research UK Gurdon Institute, The Henry Wellcome Building of Cancer and Developmental Biology, University of Cambridge CB2 1QN, UK; Department of Biochemistry, University of Cambridge, UK; Department of Genetics, University of Cambridge, UK; Department of Molecular Genetics, Erasmus University Medical Center, Dr. Molewaterplein 40, 3015GD Rotterdam, The Netherlands

## Abstract

During metazoan development, the cell cycle is remodelled to coordinate proliferation with differentiation. Developmental cues cause dramatic changes in the number and timing of replication initiation events, but the mechanisms and physiological importance of such changes are poorly understood. Cyclin-dependent kinase (CDK) is important for regulating S-phase length in many metazoa, and here we show in the nematode *Caenorhabditis elegans* that an essential function of CDK during early embryogenesis is to regulate the interactions between three replication initiation factors SLD-3, SLD-2 and MUS-101 (Dpb11/TopBP1). Mutations that bypass the requirement for CDK to generate interactions between these factors is sufficient for viability in the absence of CyclinE/Cdk2, demonstrating that this is a critical embryonic function of this cyclin/CDK complex. Both SLD-2 and SLD-3 are asymmetrically localised in the early embryo and the levels of these proteins inversely correlate with S-phase length. We also show that SLD-2 asymmetry is determined by direct interaction with the polarity protein PKC-3. This study explains the essential function of CDK for replication initiation in a metazoan and provides the first direct molecular mechanism through which polarization of the embryo is coordinated with DNA replication initiation.

**Author Summary:** How and when a cell divides changes as the cell assumes different fates. How these changes in cell division are brought about are poorly understood, but are critical to ensure that cells do not over-proliferate leading to cancer. The nematode *C. elegans* is an excellent system to study the role of cell cycle changes during animal development. Here we show that two factors SLD-2 and SLD-3 are critical to control the decision to begin genome duplication. We show that these factors are differently distributed to different cell lineages in the early embryo, which may be a key event in determining the cell cycle rate in these cells. For the first time we show that, PKC-3, a key component of the machinery that determines the front (anterior) from the back (posterior) of the embryo directly controls SLD-2 distribution, which might explain how the polarisation of the embryo causes changes in the proliferation of different cell lineages. As PKC-3 is frequently mutated in human cancers, how this factor controls cell proliferation may be important to understand tumour progression.

## Introduction

Eukaryotes replicate their genomes from multiple origins that fire throughout S-phase of the cell cycle. Programmed changes in the number, timing and position of origin firing occur during differentiation and development across many metazoa [1]. As a result, different cell types exhibit dramatic changes in the rate of S-phase and the timing with which different parts of the genome are replicated. The mechanisms and physiological importance of such changes in genome duplication during the lifetime of an organism are very poorly understood. With its highly stereotypical cell divisions, the early *C. elegans* embryo provides an ideal system to study the role of cell cycle control during development. As early as the second embryonic division, polarity cues generate cells with different S-phase lengths [2,3]. Activators of cyclin-dependent kinase (CDK) are asymmetrically distributed in the early embryo [2,4–6] and CDK activity has been shown to be important for determining the synchrony of division [6]. Despite this, how CDK controls embryonic cell cycle length is not known.

CDK plays a critical role in the initiation of DNA replication across eukaryotes [7]. In budding yeast CDK phosphorylates two essential initiation factors Sld2 and Sld3, which results in their phospho-dependent interaction with the BRCT repeats of Dpb11 [8,9]. This CDK-dependent complex results in the recruitment of additional proteins, such as the leading strand polymerase (Pol ε) and helicase activatory factors, which together allow replisome assembly by a poorly understood mechanism [10]. Phosphorylation of Sld2 and Sld3 and interaction with Dpb11 is sufficient for the function of CDK in replication initiation in yeast, as mutations that drive the interactions between these proteins can bypass the requirement for CDK to initiate replication [8,9].

Importantly, Sld3, Sld2 and Dpb11, together with another replication initiation factor Dbf4 are low abundance and rate limiting for replication initiation in yeast [11,12]. The orthologues of these same factors are also limiting for S-phase length during the early embryonic divisions in Xenopus [13]. In Drosophila, increasing CDK activity is sufficient to reduce S-phase length in the early embryo [14], although the same is not true in Xenopus or zebrafish [15,16]. It would therefore seem that limiting CDK activity and/or low levels of the key CDK substrates, Sld3 and Sld2 and their binding partners might provide a simple mechanistic explanation for how diverse organisms regulate the rate of replication initiation and thus total S-phase length. Unfortunately the testing of this hypothesis has been hampered by the difficulties in identifying the true targets of CDK in replication initiation in developmental model systems [17].

We have previously provided the first example of an essential CDK substrate required for replication initiation in a metazoan through the characterisation of *C. elegans sld-2* [18]. Mutation of the CDK sites in *sld-2* to alanine prevented the interaction with the Dpb11 orthologue MUS-101 (also known as Cut5/TopBP1) and resulted in lethality, while phospho-mimicking mutations in these CDK sites restored the interaction with MUS-101 and restored viability [18]. Having characterised *sld-2* as an essential CDK target in *C. elegans* we set out to identify and characterise the Sld3 orthologue in this organism to determine the importance of regulation of both of these substrates during development. In this study we show that CDK-dependent regulation of *sld-2* and *sld-3* is sufficient to fulfil at least in part the essential functions of cyclin E/Cdk2 in *C. elegans*. Both SLD-2 and SLD-3 are asymmetrically localised in the early embryo and the asymmetry of SLD-2 is directly regulated by an interaction with the polarity factor PKC-3. This study provides the first direct link between the cell polarity machinery and DNA replication control and pinpoints *sld-2* and *sld-3* as potentially key factors for determining S-phase length in the early embryo in *C. elegans*.

## Results

### ZK484.4 is *C. elegans* SLD-3

To identify the CDK target Sld3/Treslin in *C. elegans*, we performed homology searches using the conserved Cdc45 interaction domain (also known as the Sld3/Treslin domain, orange, Fig 1A) [19,20]. From this, we identified ZK484.4 as the best-hit for a potential orthologue of Sld3/Treslin in *C. elegans* (Fig 1A). To determine whether ZK484.4 is a functional orthologue of Sld3, we first analysed the interactions of this protein with the replication initiation factors CDC-45 and MUS-101, the *C. elegans* orthologue of Dpb11/TopBP1. Yeast two-hybrid analysis revealed that the Sld3/Treslin domain of ZK484.4 (151-388) interacted with *C. elegans* CDC-45, while the C-terminus of SLD-3 (388-873) interacted with MUS-101 (Fig 1B). Interestingly ZK484.4 interacted with the region of MUS-101 (1-448) encompassing the N-terminal BRCT repeats (Fig 1B), which is the same region of interaction between Sld3-Dpb11 in yeast and Treslin-TopBP1 in Xenopus/humans [8,9,21,22]. These conserved interactions strongly suggested that ZK484.4 is indeed the *C. elegans* orthologue of Sld3/Treslin and we hereby refer to ZK484.4 as *sld-3*. Notably we did not identify an orthologue of the Sld3/Treslin interacting protein Sld7/MTBP [23].

**Fig 1.**
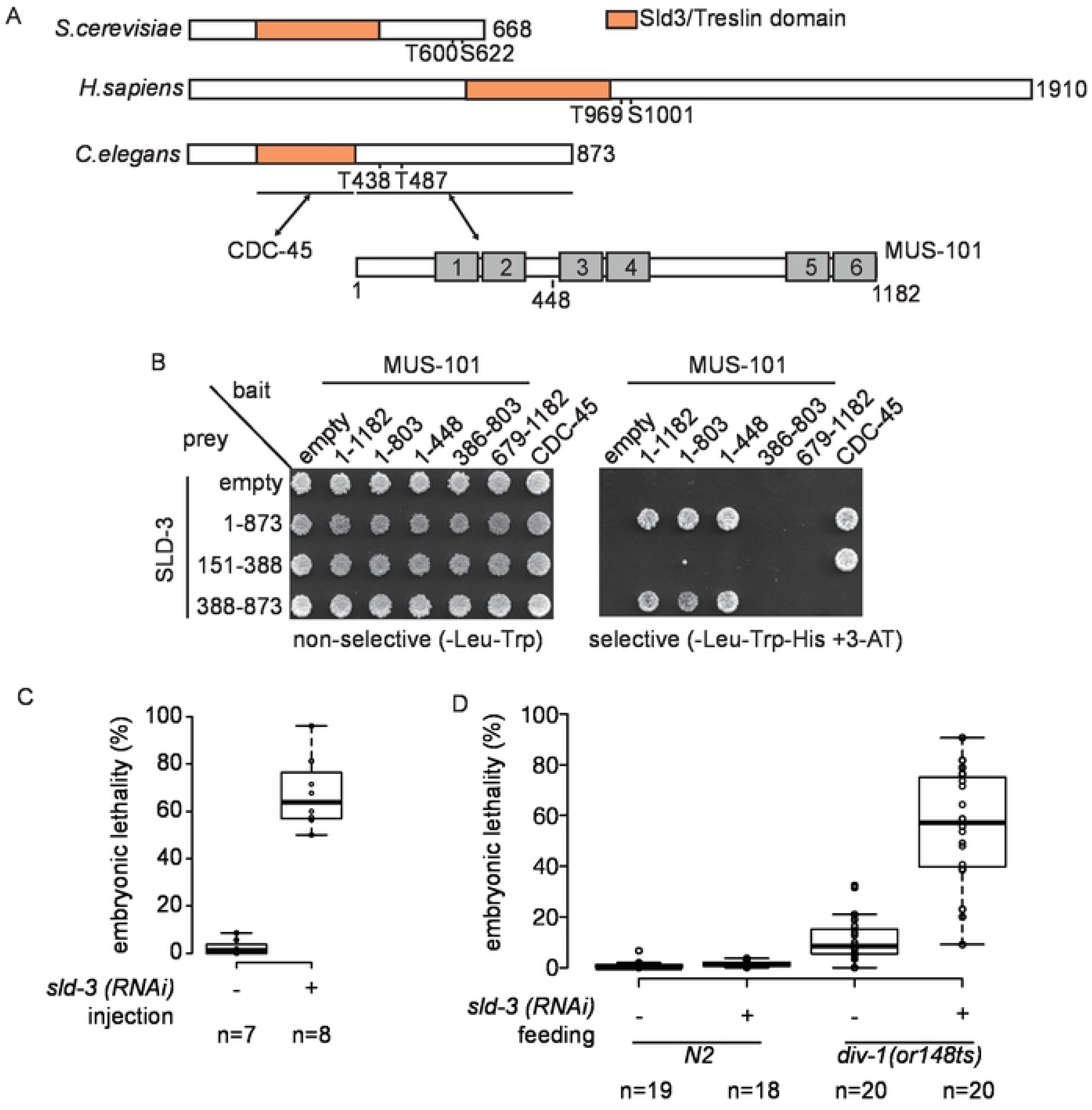
ZK484.4 is *C. elegans sld-3*. **A.** Scale diagram of Sld3, Treslin and ZK484.4 from budding yeast, humans and *C. elegans* respectively. The Cdc45 interaction (Sld3/Treslin) domain is in orange. The essential CDK sites in Sld3/Treslin and their potential orthologues in *C. elegans* are numbered. The regions of interaction between *C. elegans* ZK484.4 and CDC-45/MUS-101 are indicated below, together with a scale diagram of MUS-101 showing the 6 BRCT repeats as grey boxes. **B.** Yeast two-hybrid analysis between MUS-101 and CDC-45 bait constructs and SLD-3 prey on non-selective (-Leu-Trp) and selective medium (-Leu-Trp-His+3-AT). **C.** Box and whisker plot of embryonic lethality with and without *sld-3* RNAi by injection. **D.** As C except RNAi was performed by feeding at 21°C.

As Sld3/Treslin is essential for replication initiation across eukaryotes we set out to test whether *sld-3* is also essential in *C. elegans*. RNAi of *sld-3* by injection indeed showed that this is an essential gene (Fig 1C). Consistently with a role for *sld-3* in DNA replication, partial knock down of *sld-3* through RNAi by feeding resulted in synthetic lethality with the *div-1* mutant in the B subunit of polymerase alpha [24] at the semi-permissive temperature (Fig 1D). Together these data confirm that ZK484.4 is likely to be the functional orthologue of *sld-3*.

### *C. elegans* SLD-3 has two essential CDK sites

Sld3/Treslin is a critical CDK substrate in yeast, Xenopus extracts and human cells and CDK phosphorylation of Sld3/Treslin at two sites mediates the interaction with the N-terminal BRCT repeats of Dpb11/TopBP1 [8,9,21,25]. As *C. elegans* SLD-3 also interacts with the N-terminal region of MUS-101 (Fig 1B), we set out to determine whether CDK sites were crucial for this interaction. *C. elegans sld-3* has two CDK sites at positions 438 and 487, which show homology to the two essential sites in Sld3/Treslin (Fig 2A). Mutation of threonine 438 and 487 to alanine (hereafter called the 2A mutant) prevented the interaction between SLD-3 and MUS-101 (Fig 2B). To test whether this interaction is important *in vivo*, we inserted in the genome at a MosSCI site an additional RNAi insensitive copy of either wild type or *sld-3(2A)* fused to mCherry [26]. The expression of these MosSCI alleles was similar, as determined by mCherry fluorescence levels (see for example Fig S3B). Importantly, while the *sld-3* wild type allele fully rescued the *sld-3* RNAi lethality, the 2A mutant that cannot interact with MUS-101 could not rescue this lethality (Fig 2C). This suggests that the interaction between SLD-3 and MUS-101 is critical *in vivo* in *C. elegans*.

**Fig 2.**
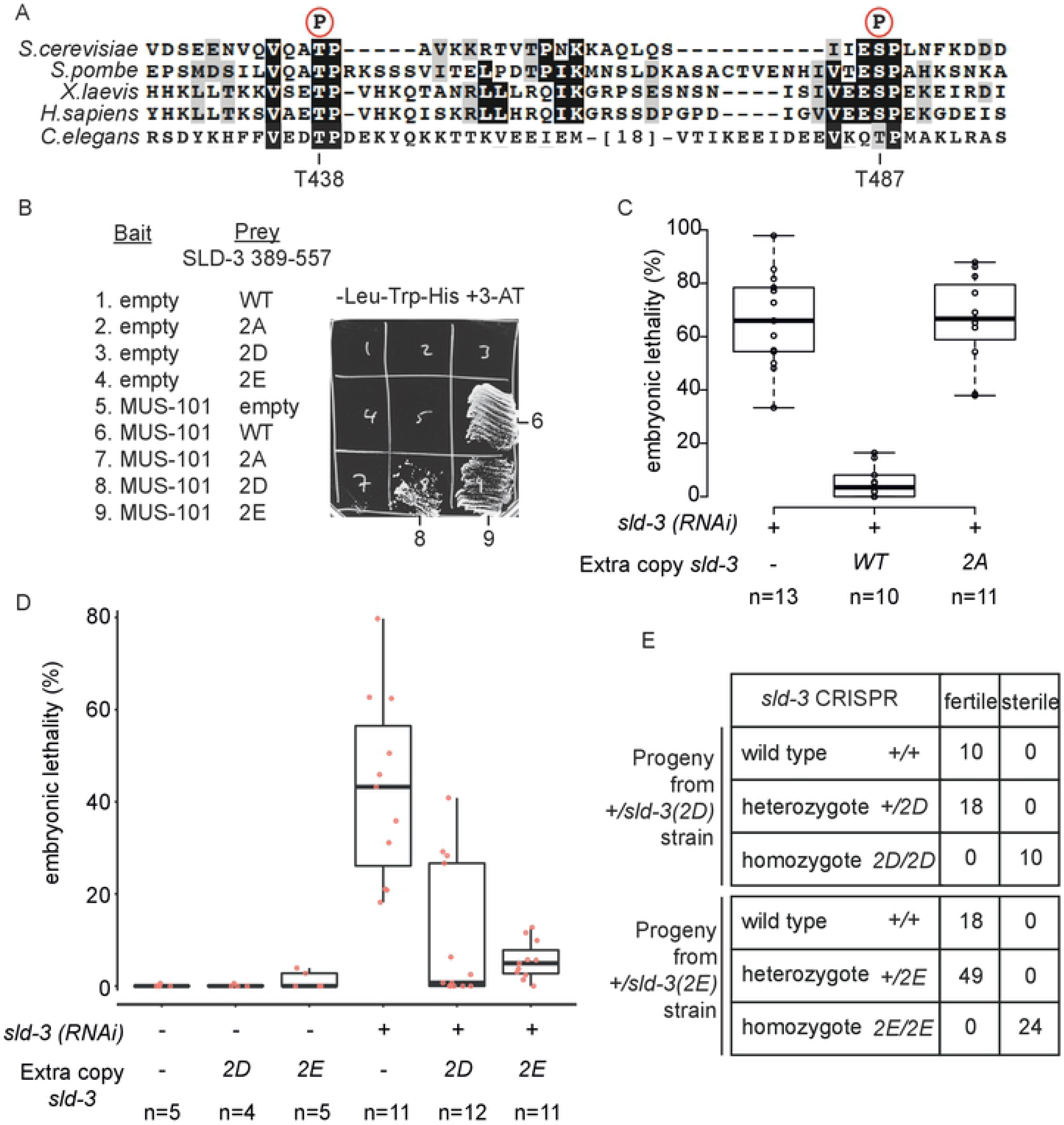
Two CDK sites in SLD-3 are essential for the interaction with MUS-101. **A.** Alignment of the CDK sites in Sld3/Treslin required for the interactions with Dpb11/TopBP1. The amino acid numbers of the two orthologous CDK sites in *C. elegans* SLD-3 are indicated below. **B.** Yeast two-hybrid analysis between MUS-101 (1-448) and SLD-3 (389-557) wild type (WT) or with the two CDK sites threonine 438 and 487 mutated to alanine (2A), aspartic acid (2D) or glutamic acid (2E). **C** and **D.** Box and whisker plot of embryonic lethality after *sld-3* RNAi by injection as in Fig 1C. The extra, RNAi insensitive copies of *sld-3* are inserted at a MosSCI site and expressed from the *mex-5* promoter. **E.** Ratio of progeny from heterozygous +/*sld-3(2D)* (top) or +/*sld-3(2E)* (bottom) parents. These 2D/2E mutations were generated by CRISPR at the endogenous *sld-3* locus.

As phosphorylation of Sld3/Treslin is required for the interaction with Dpb11/TopBP1 [8,9,21,25] we wondered whether this was also the case in *C. elegans*. Significantly, mutation of the two essential CDK sites to aspartic acid (2D) or glutamic acid (2E), which potentially mimics phosphorylation of these sites, restored the interactions with MUS-101 (Fig 2B). Critically, while *sld-3* RNAi resulted in high levels of lethality as previously shown, an RNAi insensitive copy of *sld-3(2D)* partially rescued this lethality, while the *sld-3(2E)* allele almost fully rescued the loss of wild type *sld-3* (Fig 2D), unlike the situation for human Treslin [27]. Together this shows that mutations that mimic phosphorylation of *sld-3* at these two essential sites allow MUS-101 interaction and restore viability *in vivo*.

Since expression of the *sld-3(2D)* or *(2E)* mutants as a second copy restored viability after *sld-3* RNAi, we set out to generate these alleles at the endogenous locus by CRISPR. While heterozygotes of the CRISPR-generated *sld-3(2D)* and *(2E)* mutants were viable, the homozygotes were sterile (Fig 2E). We wondered whether instead of mutating both CDK sites, mutation of just one site might be sufficient to generate viable alleles. Strains that were homozygous for either T438 or T487 mutated to alanine resulted in intermediate levels of embryonic death and infertility (Fig S1). While mutation of these individual sites to aspartic acid did not rescue the lethality/sterility, mutation of these sites to glutamic acid significantly reduced the lethality exhibited by the alanine mutants (Fig S1). Together with the analysis of the *sld-3* alleles as a second copy (Fig 2D), these data show that while alanine mutants of either of the two, or both CDK sites show lethality, phospho-mimicking mutants can bypass and rescue to some extent this lethality *in vivo*.

We are not sure why the *sld-3(2E)* allele shows high levels of viability after *sld-3* RNAi (Fig 2D), but not as a homozygous allele at the endogenous locus (Fig 2E). One possibility is that constitutive phospho-mimicking of these sites generates phenotypic issues by itself, indeed the CDK bypass mutants of Sld3 alone are sick in yeast [8]. We consider it more likely however that these alleles are simply not fully penetrant in mimicking the essential functions of *sld-3* and therefore behave as hypomorphs, which are viable in the presence of some background level of wild type protein (e.g after RNAi). Despite this, the phospho-mimicking mutants of the CDK sites in *sld-3*, which allow MUS-101 interaction, show dramatic rescue of *sld-3* RNAi lethality (Fig 2D) *in vivo*.

### Bypass of CDK site phosphorylation in SLD-3 and SLD-2 is partially sufficient for cyclin E/CDK2 function

In yeast, phospho-mimicking mutants of Sld2 and Sld3 fulfil the essential functions of CDK in DNA replication initiation [8,9]. In a previous study we characterised the Sld2 orthologue in *C. elegans* and identified the *sld-2(8D)* mutant as capable of bypassing the requirement for the CDK sites in *sld-2* to allow interaction with the C-terminus of MUS-101 [18]. Here we have identified the *sld-3(2D)* and *(2E)* mutants that can bypass the requirement for the CDK sites to generate the crucial interaction between SLD-3 and the N-terminus of MUS-101 (Fig 2). Therefore we wondered to what extent the combination of these bypass mutants of *sld-3* and *sld-2* might be able to fulfil essential functions of CDK in *C. elegans* (Fig 3A). As in yeast, we might expect such CDK bypass mutants to be dominant, which is indeed the case for *sld-2(8D)* [18]. Combination of the CDK bypass mutant *sld-2(8D)* with *sld-3(2D)* or *(2E)*, expressed as extra copies at MosSCI sites, resulted in wild type levels of fertility and viability (Fig 3B and S2A).

**Fig 3.**
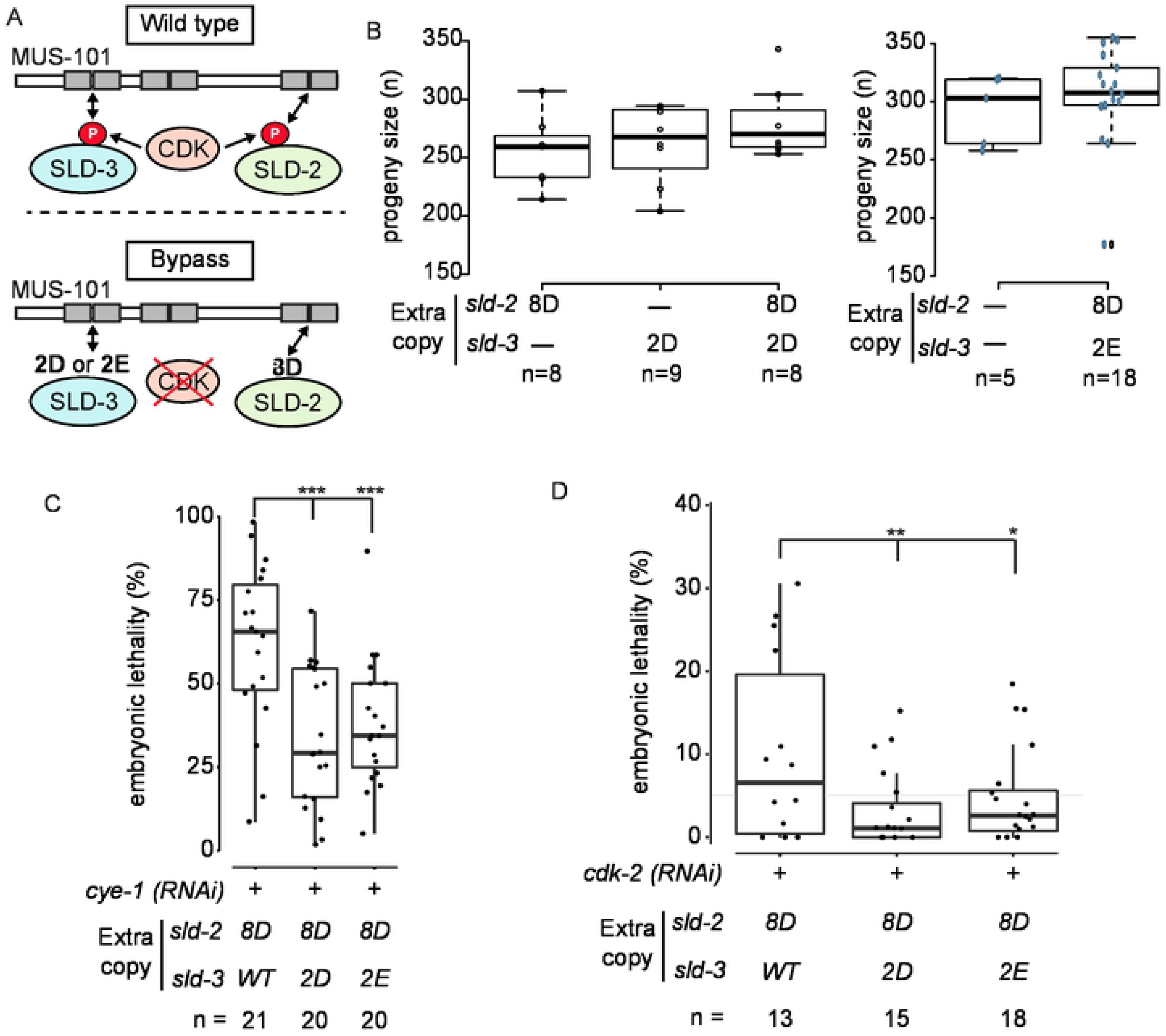
Bypass of CDK site phosphorylation in SLD-3 and SLD-2 is partially sufficient for cyclinE/CDK-2 function. **A.** CDK drives the interactions between SLD-2, SLD-3 and MUS-101 (top). The requirement for CDK can be bypassed using phospho-mimicking mutants in the essential CDK sites of SLD-2 (8D) or SLD-3 (2D or 2E). **B.** Box plot of progeny size for the indicated strains containing *sld-2* and/or *sld-3* mutants as extra copies as MosSCI insertions. **C** and **D.** Box plot of embryo lethality as in Fig 1C after *cye-1* (C) or *cdk-2* (D) RNAi by feeding in the indicated strains. *** P-value <0.005, **0.006, *0.0453.

Cyclin E/CDK2 is required for the G1-S transition and is responsible for DNA replication initiation, particularly in early embryonic divisions such as in Drosophila and Xenopus [28,29]. RNAi of either Cyclin E (*cye-1*) or *cdk-2* resulted in embryonic lethality, as expected [30] (Fig 3C-D and S2B). Expression of the *sld-2* or *sld-3* bypass alleles alone did not restore viability after *cye-1* or *cdk-2* RNAi (Fig S2B-C). Importantly combination of both *sld-2(8D)* and *sld-3(2D)* or *(2E)* resulted in significant rescue of viability of both *cye-1* and *cdk-2* RNAi (Fig 3C-D). These phenotypic rescues by the *sld-2/sld-3* bypass mutants were specific to cyclinE/Cdk2 RNAi, as we did not observe any rescue with Cyclin B1 (*cyb-1*) or Cyclin B3 (*cyb-3*) RNAi (Fig S2D-E). Unfortunately RNAi of *cya-1* (Cyclin A) had no phenotype in our hands (data not shown). Together these data show that *sld-2* and *sld-3* mutants that can bypass the requirement for the critical CDK sites for generating interactions with MUS-101 can fulfil some of the essential functions of Cyclin E/Cdk2 *in vivo* in *C. elegans*.

### SLD-2 and SLD-3 are asymmetrically localised in the early embryo

During the second embryonic division in *C. elegans*, the anterior AB cell has a faster cell cycle than the posterior P1 cell, which is in part due to a shorter S-phase in the AB cell [2]. CDK activity is potentially differentially activated in these two cells due to the asymmetric distribution of CDK regulators, such as *cdc-25* and the cyclin *cyb-3* [4,6]. We wondered to what extent SLD-2 and SLD-3 regulation by CDK might contribute to this asynchrony of cell division, so we analysed the AB/P1 cell cycle duration using the *sld-2/sld-3* CDK bypass alleles. Fig 4A shows that the duration of the AB and P1 divisions remained very similar in the *sld-2(8D)/sld-2(2E)* mutant relative to wild type, suggesting that CDK phosphorylation of these targets alone is not limiting for S-phase duration in either of these cell divisions.

**Fig 4.**
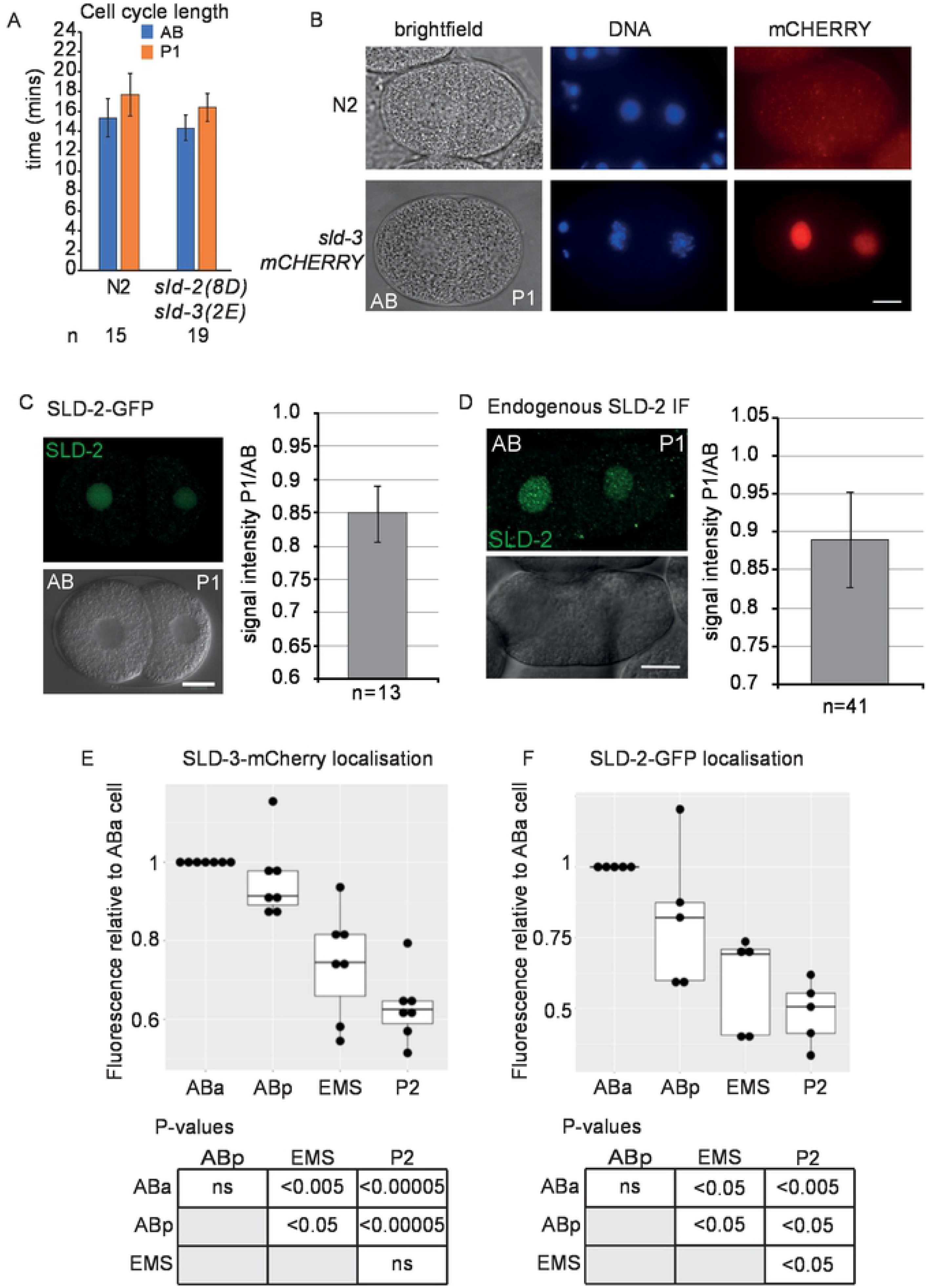
SLD-3 and SLD-2 are asymmetrically localised in the early embryo Cell. **A.** cycle length in the AB and P1 cell of the two-cell embryo in the indicated strains. Error bars are SD. **B.** Images of two-cell embryos of wild type (N2) or containing *mex-5p::sld-3-mCherry::tbb-2 3-UTR* construct integrated at a MosSCI site. mCherry was visualised by IF, DNA was stained with Hoechst. Scale bar is 10μm **C.** As B. Images of two-cell embryo containing *mex-5p::sld-2-gfp::tbb-2 3UTR* integrated at a MosSCI site. GFP signal was detected with Confocal Laser Scanning Microscopy. Lower Image shows DIC channel. Scale bar is 10 μm. (Right) Graph indicates the ratio of GFP signal intensity from the P1 cell over the signal from the AB cell. Error bars are 95% CI. **D.** As C, except endogenous SLD-2 was detected by immunofluorescence. **E** and **F.** Quantitation of SLD-3-mCherry signal by IF (E) and SLD-2-GFP signal by fluorescence imaging (F) in the four-cell embryo. The signal in the ABa cell was set to 1. n=7 for each measurement for E and n=5 for F. Below the p-values were obtained by paired t-tests, ns is not significant.

During this analysis of AB/P1 cycle length using the MosSCI *sld-3* and *sld-2* alleles, which are tagged with mCherry and GFP respectively, we observed that both SLD-3 and SLD-2 showed asymmetric localisation, with more protein in the AB cell nucleus, than P1 (Fig 4B/4C). This asymmetry was not limited to the MosSCI alleles, as we obtained a similar result using immuno-fluorescence of endogenous SLD-2 (Fig 4D). Interestingly the presence or absence of the essential CDK sites did not affect the asymmetric localisation of SLD-3 (Fig S3A-B). The asymmetry we observed was not an artefact of embryo staging, as the difference in abundance of SLD-2 was detected throughout interphase (Fig S3C).

Asymmetric and asynchronous divisions continue beyond the 2-cell stage, with the descendants of the AB cell (ABa and Abp) having shorter cell cycles than the descendants of the P1 cell (EMS and P2) with P2 having the longest S-phase of these cells [3,31]. We analysed the abundance of SLD-2 and SLD-3 in 4-cell stage embryos and these two proteins remained asymmetric at this stage with EMS and P2 having significantly less protein than the AB cell lineage (Fig 4E-F). SLD-2 abundance was also significantly lower in the P2 cell than the EMS cell (Fig 4F). Together these data show that SLD-2 and SLD-3 are present at levels that inversely correlate with S-phase length in the 2- and 4-cell *C. elegans* embryo.

### PAR proteins control SLD-2 asymmetry

The PAR polarity proteins (PAR-1 to -6) and PKC-3, which specify the anterior-posterior (A–P) axis in the early embryo, also regulate the asynchrony of cell division between the AB and P1 blastomeres [2]. *par-3* and *pkc-3* mutants divide synchronously and symmetrically at the 2-cell stage [2,32] and significantly loss of function of either of these polarity genes resulted in subsequent symmetrical localisation of SLD-2 in the AB and P1 cell (Fig 5A-5B).

**Fig 5.**
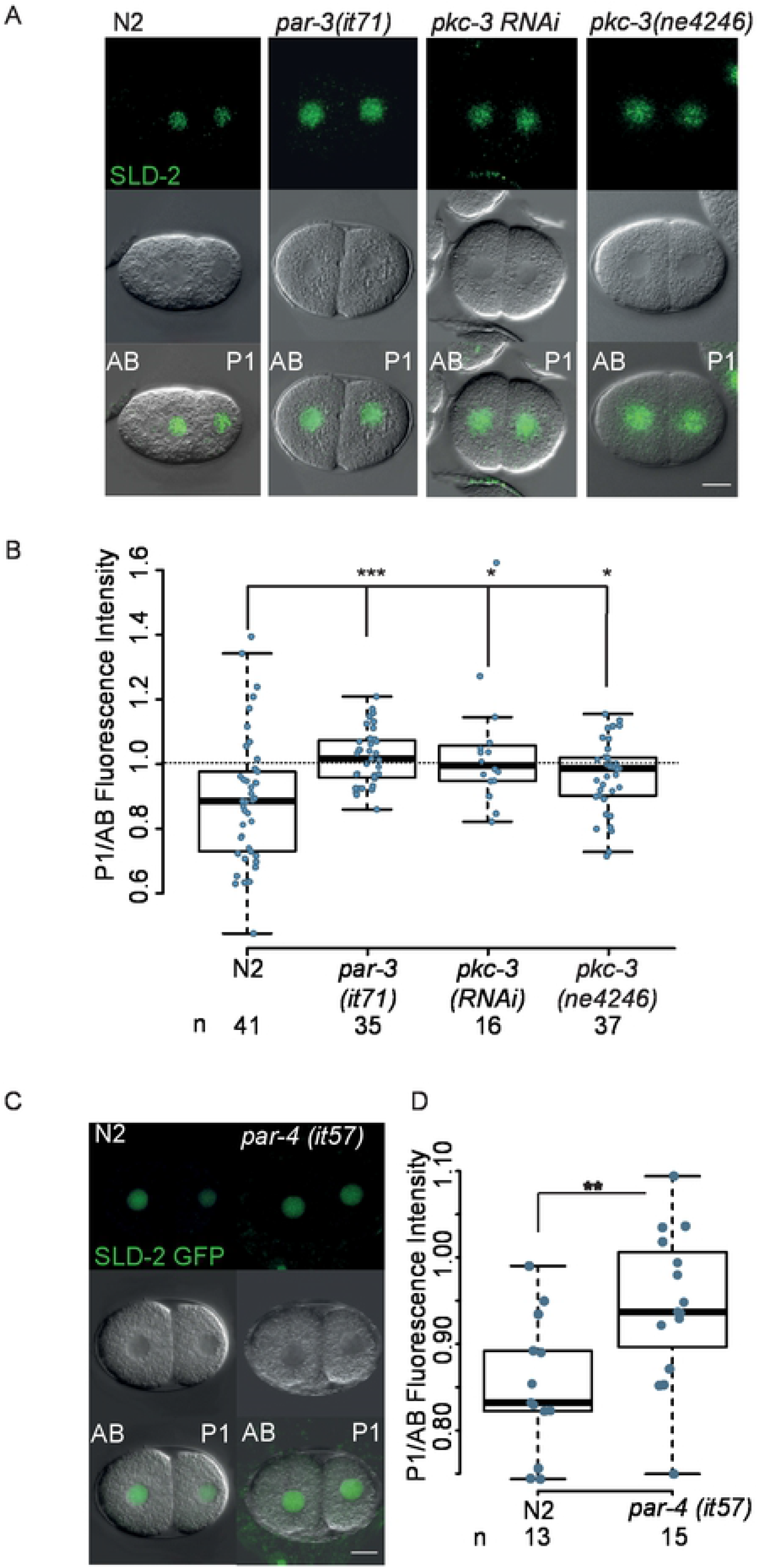
SLD-2 asymmetry is polarity-dependent. **A.** Images of two-cell embryos from the indicated strains after SLD-2 IF. Scale bar is 10 μm. N2, *par-3(it71)* and *pkc-3 RNAi* animals were grown at 20°C. The temperature-sensitive strain *pkc-3(ne4246)* was grown at 25°C **B.** Box plot of the data from A. p-values were obtained using the Wilcoxon rank sum test. **C.** Images of two-cell embryo of wild type (N2) and the temperature-sensitive *par-4(it57)* mutant containing *mex-5p::sld-2-gfp::tbb-2 3UTR* grown at 25°C. The fluorescent GFP signal was detected with Laser Confocal Scanning Microscopy. Scale bar is 10 μm. **D.** Box plot of the data from C. p-values were obtained using the Wilcoxon rank sum test.

We have previously shown in Xenopus that nuclear-to-cytoplasmic ratios can affect S-phase length due to the amount of limiting replication initiation factors inherited after cell division [13]. As *par-3* and *pkc-3* mutants divide symmetrically (Fig 5A), we wondered whether the subsequent symmetry of SLD-2 was simply a consequence of equal distribution of cellular content after division. To test this we analysed the distribution of SLD-2 in *par-4* mutant embryos, which divide synchronously but still asymmetrically at the 2-cell stage, resulting in AB/P1 cells of similar size to wild type [33]. Significantly, SLD-2 was symmetrically localised in *par-4* mutant embryos, even though the P1 cell is smaller than the AB cell in these mutants (Fig 5C-D). Together this suggests that SLD-2 localisation is actively regulated by the PAR protein network not simply by the cellular volume at division.

### PKC-3 interacts with SLD-2 and causes SLD-2 asymmetry in the embryo

To understand the molecular mechanism of SLD-2 asymmetry, we performed a yeast two-hybrid screen between SLD-2 and a cDNA library from *C. elegans* embryos (data not shown). One of the hits from this screen was the polarity factor *pkc-3*, which is essential for defining the anterior domain in the 1-cell embryo [34]. SLD-2 interacts with the PKC-3 region 94-184, which encompasses the pseudosubstrate (PS) and C1 domains (Fig 6A). To assess the function of this interaction *in vivo*, we set out to identify a separation of function mutant in *sld-2*, which lacked the PKC-3 interaction. Using yeast two-hybrid analysis we narrowed down the interaction to the very C-terminus of SLD-2, region 232-249 (Fig 6B). This is a highly basic region of SLD-2 (Fig S4A), which lacks any CDK sites. Indeed the SLD-2 mutant lacking all 8 CDK sites (either mutated to alanine or aspartic acid, 8A/8D) still interacted with PKC-3 (Fig 6B). To identify a mutant that no longer interacted with PKC-3 we made scanning mutations in the region 232-249 (Fig S4B-C). A mutation that converted the very C-terminal 4 amino acids from KKKY to the acidic residues EDDD indeed resulted in loss of the interaction with PKC-3 (Fig 6B and S4B-C) and we hereafter refer to this mutant as *sld-2(EDDD)*. To check whether these mutations affect the essential functions of *sld-2*, we tested whether *sld-2(EDDD)* expression rescued the lethality of *sld-2* RNAi. Insertion of either *sld-2* wild type or the *EDDD* mutant at a MosSCI site fully rescued the lethality of *sld-2* RNAi (Fig 6C) strongly suggesting that the *sld-2(EDDD)* mutant is not defective in any of the essential functions of *sld-2*.

**Fig 6.**
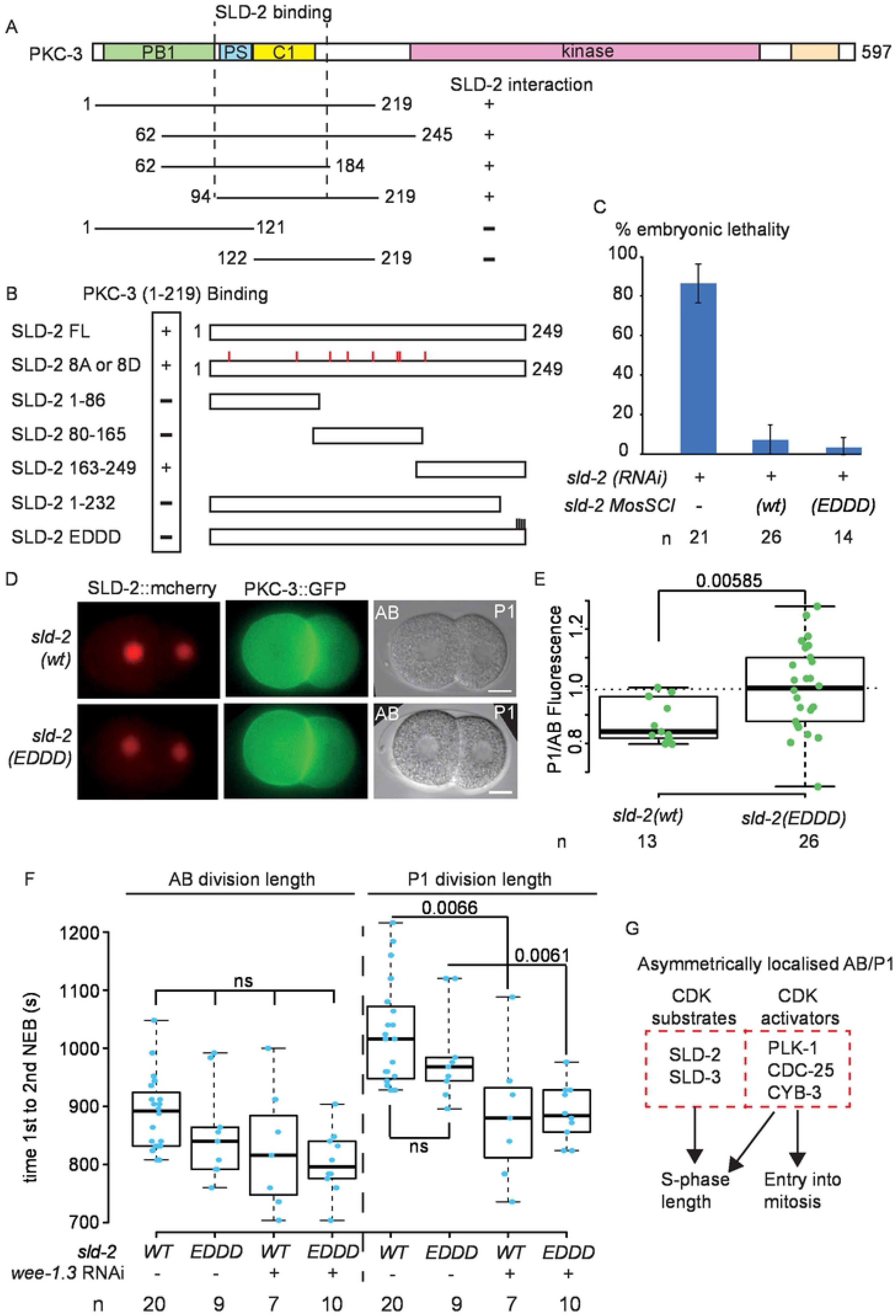
SLD-2 asymmetry is PKC-3 interaction-dependent. **A.** (Top) Scale diagram of *C. elegans* PKC-3. (Bottom) Diagram of PKC-3 fragments that were tested for yeast two-hybrid interaction with full length SLD-2. +/− represents whether the interaction was positive or negative. **B.** Scale diagram of fragments of SLD-2 that were tested for yeast two-hybrid interaction with PKC-3 (1-219). +/− represents whether the interaction was positive or negative. SLD-2 8A or 8D refers to all 8 CDK sites mutated either alanine or aspartic acid. SLD-2 EDDD refers to residues 246-249 (KKKY) mutated to EDDD. **C.** Embryonic lethality 16-40hrs post-injection from the wild type strain (N2) or MosSCI generated *mex-5p:::sld-2(WT)::gfp::tbb-2 3UTR* or *mex-5p::sld-2(EDDD)::gfp::tbb-2 3UTR* strains. SLD-2 RNAi was done by injection. Error bars are 95% CI. **D.** Live imaging of PKC-3 GFP and SLD-2 mCherry or SLD-2 EDDD mCherry generated by CRISPR. Scale bar is 10 μm. **E.** Detection of the nuclear SLD-2 signal by IF. Box plot graph shows the ratio of the fluorescent signal intensity from the P1 cell over the signal from the AB cell from the wild type strain (N2) and the mutant *sld-2 EDDD* generated by CRISPR. **F.** Box plot of the AB and P1 cell cycle length from the indicated strains with and without *wee1.3* RNAi by feeding. Cell cycle length was calculated as the time from the P0 cell nuclear envelope breakdown (NEB) to the AB or P1 NEB from 8s time lapse movies. p-values were calculated using the Wilcoxon test. **G.** Both CDK substrates (SLD-2 and SLD-3) and CDK activators (PLK-1, CDC-25 and CYB-3) are asymmetrically localised at the 2-cell stage in *C. elegans*. The relative contribution of these localisations to asynchronous embryonic cell cycle lengths remains to be determined.

To investigate the significance of the SLD-2 interaction with PKC-3 for SLD-2 localisation we generated *sld-2(WT)* and *sld-2(EDDD)* alleles by CRISPR. Homozygous *sld-2(EDDD)* strains were viable and showed no sterility phenotypes as expected from the MosSCI strains (Fig 6C and data not shown). Importantly while the wild type SLD-2 showed asymmetric localisation in the AB cell versus the P1 cell in 2-cell embryos as expected, the *sld-2(EDDD)* mutant which can no longer interact with PKC-3 exhibited equal localisation in AB and P1 cells (Fig 6D and 6E). This suggested that the interaction of SLD-2 with PKC-3 is important for the asymmetric localisation of SLD-2 in the early *C. elegans* embryo. Although PKC-3 is largely cytoplasmic and SLD-2 is mostly nuclear, SLD-2 becomes entirely cytoplasmic upon nuclear envelope breakdown and we do observe an enrichment of both nuclear and cytoplasmic SLD-2, in the AB versus the P1 cell (Fig S4D).

Having identified a mutant of *sld-2* that is no longer asymmetrically localised in 2-cell embryos, we wondered if this had an effect on the cell cycle duration of this stage. The *sld-2(EDDD)* mutant alone had no effect on the duration of the AB or P1 cycle length (Fig 6F). Previous studies have shown the importance of the inhibitory phosphorylation of CDK for elongating the P1 cell division cycle [6] and we also observed a significant reduction in P1 cell cycle length after RNAi of the CDK inhibitory kinase *wee-1* (Fig 6F). Combined inhibition of *wee-1* with the *sld-2(EDDD)* mutant did not further reduce the P1 or AB cell cycle lengths (Fig 6F). Together these data show that the PAR protein network controls SLD-2 asymmetry through PKC-3 interaction, but on it’s own symmetrical localisation of SLD-2 is not sufficient to advance the cell cycle at the 2-cell stage.

## Discussion

It is vital for all organisms to make a perfect copy of the genome in every cell division. For eukaryotes this is achieved in large part by linking DNA replication control to CDK activation at the G1-S transition [7]. CDK plays a vital dual role in DNA replication, both as an inhibitor of the helicase loading step in the initiation reaction (a process called licensing) and as an activator of these loaded helicases during replisome assembly. In budding yeast, CDK activates replisome assembly by phosphorylation of Sld2 and Sld3, but the relative contribution of phosphorylation of these two proteins to replication initiation differs in other species [17]. CDK phosphorylation of the metazoan orthologue of Sld3 (Treslin/Ticrr/C15orf42) has been shown to be important for S-phase progression in human cells in culture [21,25,27], but evidence for an essential role for Sld2 (RecQ4/RecQL4) phosphorylation in vertebrate cells is lacking. Conversely, CDK phosphorylation of Sld3 is not essential in the fission yeast *S. pombe* and Sld3 orthologues are so far absent in *D. melanogaster* [17].

By characterising *sld-2* and *sld-3* in the nematode *C. elegans*, we show for the first time outside of budding yeast that both of these proteins mediate essential interactions with MUS-101 (Dpb11/Cut5/TopBP1) through critical CDK sites (Fig 1–2 and [18]). Importantly phospho-mimicking mutants in both *sld-2* and *sld-3* that drive interactions with MUS-101 are partially sufficient for cyclin E/Cdk2 function in *C. elegans* (Fig 3). As the rescue of the *cye-1/cdk-2* RNAi with *sld-2(8D)/sld-3(8E)* is only partial we cannot rule out that there may be other CDK targets required for replication initiation in *C. elegans*, although it is also the case that the D and E mutants of *sld-2/sld-3* are not perfect phospho-mimics (Fig 2D). In addition, Cyclin E/Cdk2 has multiple functions in *C. elegans* such as contributing to embryo polarity [35] and cell cycle re-entry of differentiated cells [30,36]. While it is possible that some functions of cyclin E/Cdk2 may be compensated by other CDK complexes after *cye-1/cdk-2* RNAi, it is important to note that, apart from a small number of blast cells, all cells differentiate and become post-mitotic before the completion of embryonic development in *C. elegans* [31]. Therefore our viability assays only assess the contribution of CyclinE/Cdk2 to cell proliferation during early embryogenesis.

A surprising feature of the *sld-2(8D)/sld-3(8E)* double mutant strain is that it is perfectly viable and fertile (Fig 2B and S2). Switch like activation of CDK at the G1-S transition is required to completely separate the period of replication licensing from initiation and bypass of phosphorylation of Sld2 and Sld3 results in genome instability and death in yeast [8,37]. Additional layers of regulation may contribute to viability in the *sld-2(8D)/sld-3(8E)* strain and one possibility is the activity of the Dbf4-dependent kinase DDK. Like CDK, DDK is important for replication initiation and Dbf4 is degraded by the APC/C in G1 phase in yeast, helping to prevent precocious replication initiation [7]. We have previously shown that APC/C-dependent control of Dbf4 is important in strains that mimic phosphorylation of Sld2/3 in yeast [8]. How the *sld-2(8D)/sld-3(8E)* strain maintains a separation of licensing from initiation in *C. elegans* remains to be determined.

The early embryonic divisions in many metazoa, such as in Drosophila, zebrafish and Xenopus, are extremely rapid, lack gap phases and are characterised by high rates of replication initiation. Cell cycle lengthening in these embryonic divisions coincides with activation of DNA damage checkpoint kinases and the down regulation of cyclin-dependent kinase (CDK) activity, through the inhibitory phosphorylation of CDK by Wee1 and down-regulation of the counteracting phosphatase Cdc25 (String/Twine in Drosophila) [38,39]. Inhibitory phosphorylation of CDK is likely critical for the introduction of G2 phase and for delaying entry into mitosis. In Drosophila however increasing CDK activity can also reduce S-phase length at the mid-blastula transition (MBT) [14], although expression of CDK mutants that cannot be inhibited by Wee1 does not affect S-phase length at the MBT in Xenopus or zebrafish [15,16].

In *C. elegans*, CDC-25 and the Polo-like kinase PLK-1 (which increases the nuclear accumulation of CDC-25) preferentially localise to the faster dividing AB cell in the early embryo [2,4–6], while checkpoint activation has been proposed to preferentially occur in the P1 cell [40]. RNAi of *wee-1* in *C. elegans* indeed results in faster division of the P1 cell [6]. Therefore in both Drosophila and *C. elegans*, inhibitory phosphorylation of CDK plays an important role in cell cycle lengthening in the embryo. Despite this, in both of these organisms cell cycle elongation begins with changes in replication initiation [41,42], but how this is achieved is not clear. In Drosophila embryos, CDK activity prevents the chromatin binding of Rif1, and loss of Rif1 to a large extent prevents normal cell cycle elongation in cycle 14 [43]. Rif1 is known to inhibit replication initiation through counteraction of DDK, but also causes changes in chromatin structure [44]. Despite this, RNAi of Rif1 is not sufficient to accelerate the early embryonic divisions in *C. elegans* (MK and PZ data not shown).

Here we show that bypass of SLD-2 and SLD-3 activation by CDK is not sufficient to change the cell cycle length in the early embryo (Fig 4A). This suggests that CDK phosphorylation of these two replication substrates is not limiting for S-phase length at least at the two-cell stage. Instead we show that the SLD-2 and SLD-3 proteins themselves are asymmetrically distributed (Fig 4–6). It is striking that both the regulators and the substrates of CDK are asymmetrically localised in the AB versus P1 cell in *C. elegans* (Fig 6G and [2,4–6]). Although symmetric localisation of SLD-2 alone was not sufficient for alter the early embryonic divisions (Fig 6F), we do not currently know how SLD-3 asymmetry is controlled to test the effect of equal distribution of both proteins towards cell cycle length.

SLD-2 asymmetry in the *C. elegans* embryo is controlled by direct interaction with the polarity factor PKC-3 (Fig 6), which is preferentially localised at the anterior of the embryo [32,45]. A simple mechanistic explanation for SLD-2 accumulation in the anterior AB nucleus over the posterior P1 nucleus is therefore that SLD-2 becomes enriched in the AB cytoplasm (Fig S4D) by virtue of the established asymmetry of PKC-3. In line with this hypothesis, the localisation of both SLD-2 and the anterior polarity proteins, including PKC-3 are dependent on *par-3* and *par-4* (Fig 4 and [46,47]). Although cell polarity has been shown to be required for S-phase length control in the early *C. elegans* embryo [41], to our knowledge we have provided the first direct link between the polarity network proteins and factors that are essential for DNA replication initiation. This study may provide a platform to understand the mechanism by which programmed developmental cues directly influence S-phase length. As the human *pkc-3* orthologues are frequently mutated in cancers [48], this new link between atypical PKC and factors required for genome duplication may provide a novel mechanism by which this tumour suppressor affects cell proliferation.

## Materials and Methods

### Strains

Standard conditions were used to maintain *C. elegans* cultures (Brenner, 1974). The *C. elegans* Bristol strain N2 was used as wild type strain. Strains created by MosSCI contain wobbled versions of *sld-2* or *sld-3*. The following strains were used in this study: JA1564 (*weSi35 [Pmex-5::sld-2(wt)::egfp/tbb-2 3’UTR; cb-unc-119(*+)] II; *cb-unc-119 (ed9) III)*, JA1563*(weSi34 [Pmex-5::sld-2(8D)::egfp/tbb-2 3’UTR; cb-unc-119(*+)] II; *cb-unc-119(ed9)* III) [18], KK300 *(par-4(it57*ts*)V)* [33], KK571 *(lon-1(e185) par-3(it71)/qC1 [dpy-19(e1259) glp-1(q339)] III)* [49], KK1228 *(pkc-3(it309 [gfp::pkc-3])* II), WM150 *(pkc-3(ne4246)*II) [32] and EU548 (*div-1(or148*ts) III) [24].

Strains introduced in this study.

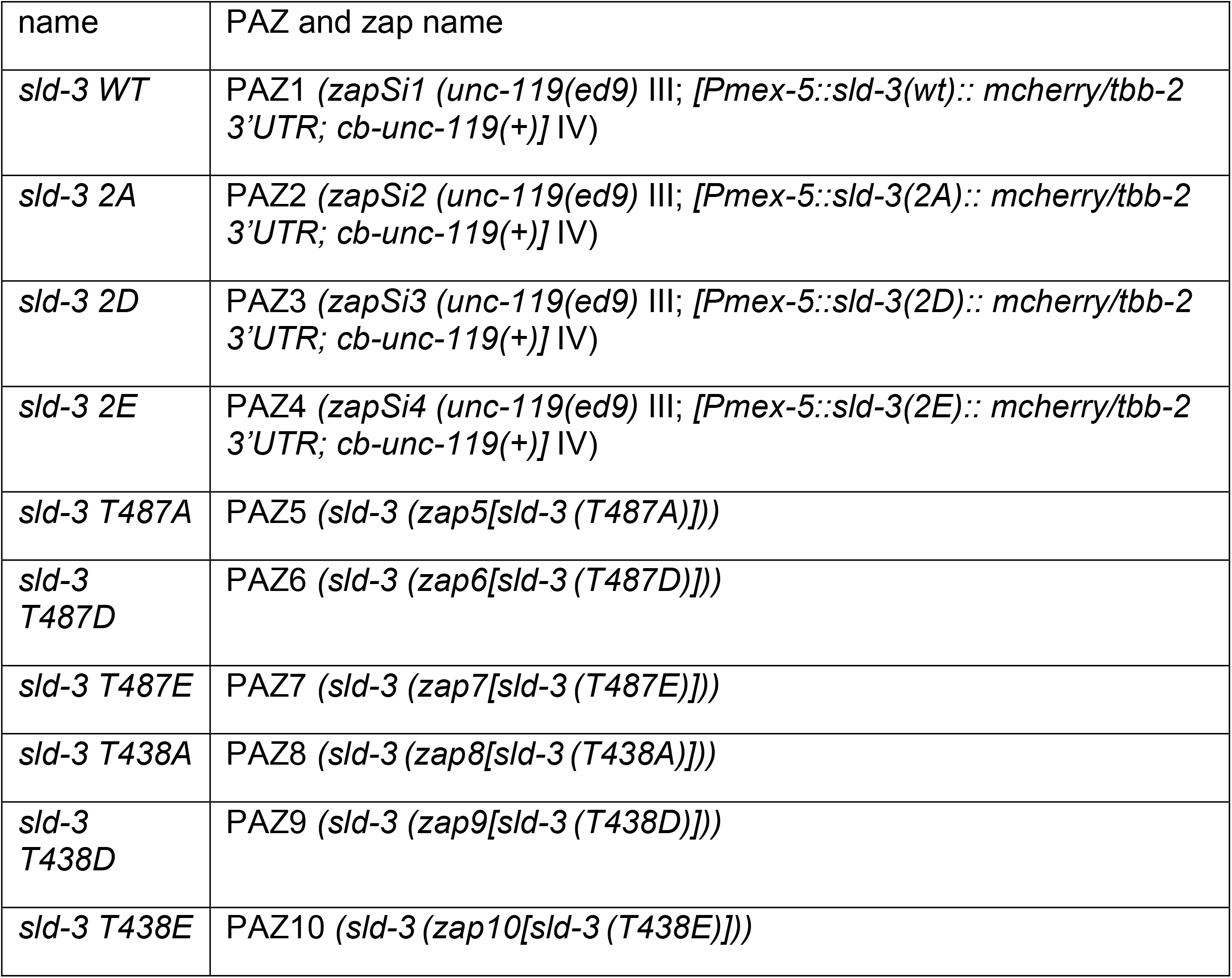
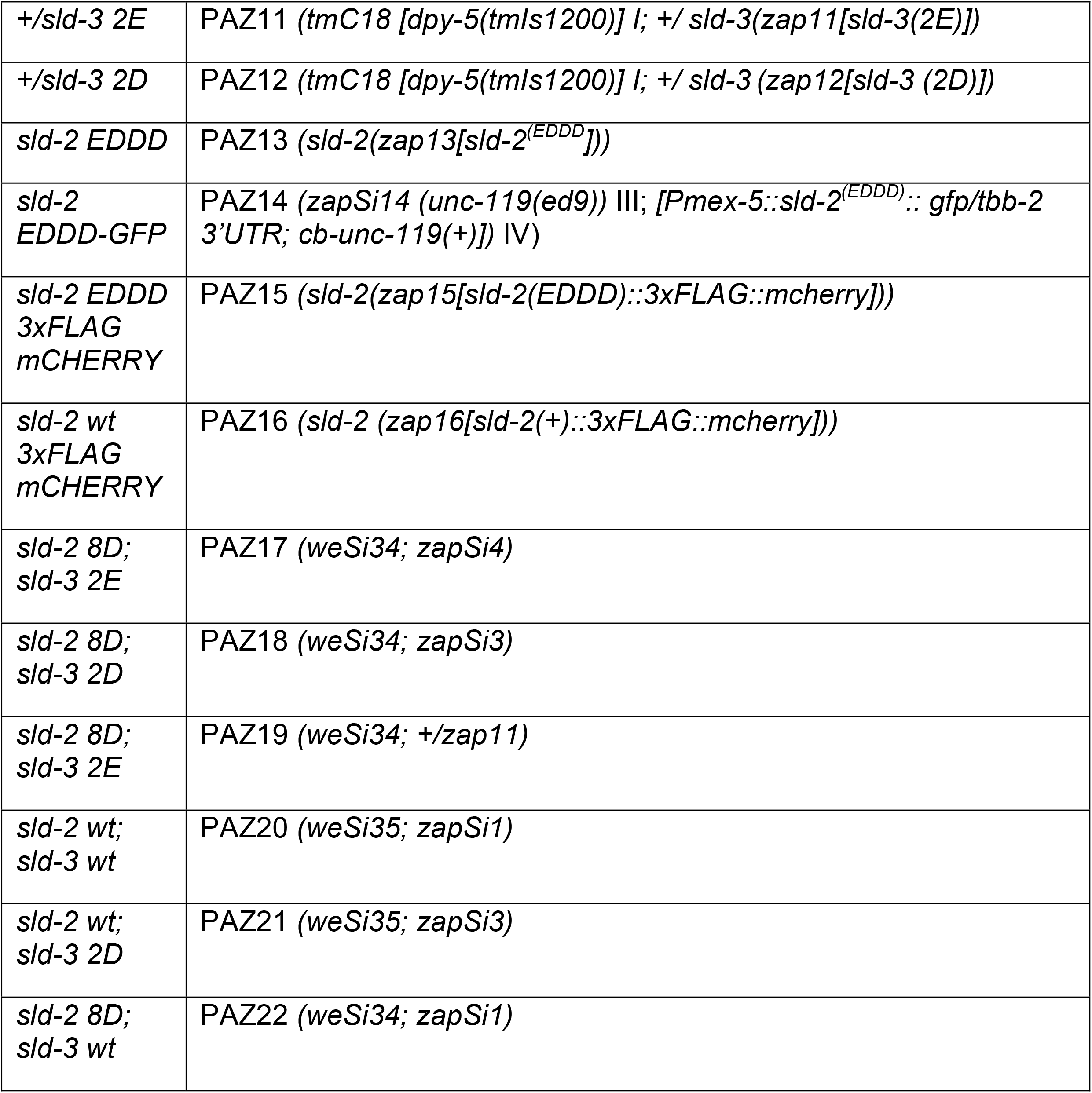

### Yeast-Two-Hybrid assays

Performed as previously described [18].

### Immunostaining

SLD-2 Immunofluorescence: Young adults were cut on a slide in a drop of M9 to release the young embryos. Embryos were freeze cracked and fixation with additional antibody incubation and washing steps were performed as described in [18]. The primary antibody was rabbit anti-SLD-2 (Ab 5058; [18]). SLD-2 antibody was used in a dilution of 1:100. Secondary antibody labelled with AlexaFluor488 anti-rabbit were obtained from Molecular Probes and used in a dilution of 1:500. Hoechst stain was added 1:1000 into the secondary antibody dilution. Vectashield® Antifading Mounting Media was used for mounting. The temperature-sensitive mutant *pkc-3(ne4246)* was kept at 25°C overnight before the immunofluorescence experiment.

SLD-3 mCHERRY Immunofluorescence: Immunofluorescence was performed using a protocol adapted from [18]. Young adult hermaphrodites were cut to release embryos onto 0.1% poly-lysine (Sigma, P8920)-coated slides. Slides were covered with a 22×50mm coverslip and frozen on dry ice for 20 minutes. The coverslip was quickly removed while the slide where still frozen to permeabilise the embryos. Embryos were fixed in ice cold methanol for 30 seconds. Slides were additionally fixed in a fixing solution containing 4% Paraformaldehyde, 80mM Hepes, 1.6mM MgSO4 and 0.8mM EGTA in PBS for 20 minutes at room temperature (RT). Samples were then washed in PBS and 0.2% Tween 20 (PBST) five times over 30 minutes, followed by blocking in 1% BSA in PBST (PBSTB) for one hour at RT. Slides were incubated in the primary antibody rabbit anti-RFP (600-401-379; Rockland antibodies and assays) (1:200) PBSTB solution overnight at 4°C. Slides were washed in PBST five times over 30 minutes, followed by incubation with the secondary antibody Alexa Fluor 594-conjugated donkey anti-rabbit antibody (A-21207; Molecular Probes) (1:500) and Hoechst 33342 stain (1:1000 final concentration 1 μg/mL) in PBSTB at RT for one hour. Samples were washed in PBST five times over 30 minutes and PBS five times over 20 minutes. Vectashield® Antifading Mounting Media was used for mounting.

### RNAi by feeding

The temperature-sensitive mutant strain EU548 (div-1(or148)) was synchronized by bleaching and grown at 15°C together with N2 wild type control. RNAi inducing plates were spotted with *sld-3* RNAi (this study, ZK484.4 ORF was cloned into L4440 plasmid) bacteria grown at 37°C for 7hrs. RNAi bacteria grown at 37°C for 7hrs. L1 worms were seeded on RNAi inducing plates and kept at 21°C until adulthood. Young adults were singled on separate NGM plates and the embryonic lethality of their progeny was determined.

For the embryonic lethality of *cdk-2*, cyb-1 and *cye-1* RNAi bacteria were grown at 37°C for 7hrs in LB containing Ampicillin. Worm strains were synchronized by bleaching. L4 worms were seeded on RNAi inducing plates with the respective RNAi bacteria until they reached adulthood and were allowed to lay eggs for 24hrs. Plates were kept at 25°C. The percentage of embryonic lethality of the F1 generation was calculated by counting the number of hatched and unhatched progeny.

The wildtype strain N2 was used for *pkc-3* RNAi. Worms were synchronized by bleaching. Plates were kept at 20°C. Mid L3 animals were seeded on RNAi inducing plates. Young adults were used for SLD-2 immunofluorescence staining. A subset of worms was singled to confirm PKC-3 knockdown by assessing embryonic lethality (data not shown).

wee1.3 RNAi by feeding was performed with the wildtype strain N2 and the Crispr mutant *sld-2(zap13[sld-2(EDDD])*. Worm strains were synchronized by bleaching. Bacteria were grown at 37°C for 7hrs in LB containing Ampicillin. A dilution of 10% (v/v) wee-1.3 bacteria in L4440 control bacteria was used for seeding RNAi plates. Because wee-1.3 bacteria induced sterility, young adults were used for RNAi induction overnight in 20°C. The next day the cell division timing was assessed by time-lapse movies. A subset of worms was transferred to new plates to confirm wee-1.3 knockdown through embryonic lethality (data not shown).

cyb-3 RNAi bacteria were grown in 37°C for 7hrs in LB containing Ampicillin. 2.5% (v/v) cyb-3 bacteria diluted in L4440 control bacteria was used for seeding RNAi inducing plates. Worms were synchronized by bleaching. Plates were kept at 20°C. L3 larva were seeded on RNAi plates. Young adult worms were singled out and allowed to lay eggs for 24hrs. After additional 36hrs the embryonic lethality was determined by counting the hatched and the unhatched progeny.

All RNAi experiments included the feeding of L4440 bacteria as a control.

### RNAi by injections

*sld-3* RNAi injections: N2, PAZ1, PAZ2, PAZ3 and PAZ4 young adult hermaphrodites were injected with *sld-3* double stranded RNA, containing the entire coding region of ZK484.4 with a concentration of 100ng/μl. Injected worms were kept at 20°C and singled to separate NGM plates to assess the embryonic lethality in the F1 generation.

*sld-2* RNAi injections: N2, JA1564 and PAZ14 were synchronized by bleaching and grown at 20°C to the young adult stage. Young hermaphrodites were injected with *sld-2* double stranded RNA prepared from T12F5.1 with a concentration of 150ng/μl. Injected worms were incubated at 20°C. The injected worms were singled out after 16hrs post-injection and transferred on new plates every 24hrs for 3 days. The plates were assessed for total egg production and lethality in the embryos.

### Microscopy and image analysis

Immunofluorescence of SLD-3 mCHERRY was visualized on a DeltaVision Deconvolution microscope. Z-stack images were taken and deconvoluted using the associated software.

The fluorescent signal of the nuclei was analysed with ImageJ. Z-stack images were combined using the Z-stack tool for maximum projection. The signals of the AB and p1 nuclei were normalized against background and the signal ratio of P1/AB visualized using R.

The immunofluorescent signal of SLD-2 staining and in-vivo imaging of the SLD-2 GFP signal in JA1563 and *weSi35; par-4(it57)* were visualized using Leica S8 Laser Confocal microscope. Z-stack images through the whole embryo were taken. Images were analysed with ImageJ. Using the maximum projection z-stack tool the fluorescent signal of the nuclei was measured and normalized for background signal.

#### Relative Fluorescence analysis of SLD-3 and SLD-2 signal in the four-cell embryo

Z-stack images of SLD-3 and SLD-2 IF was analysed using ImageJ. The fluorescent intensity signal of the different nuclei was obtained using the maximum projection tool. The signal in Aba was set to 1. The signal was normalized for background using the signal of the AB cytoplasm.

### SLD-2 GFP localization kinetics

Time lapse z-stack movies of 1 min intervals were taken from early embryos of JA1564 with a Deltavision microscope using a 60× oil objective. The embryonic development of the pronuclei meeting to the nuclear envelope breakdown of the AB cell was recorded. The fluorescent signal intensity in the AB cell and the P1 cell was analysed using automated image analysis script run in ImageJ. The newly developed programme recognises the AB and the P1 cells in the Nomarski channel. Within the cells it measures an expanded area in the fluorescent GFP channel. The nuclear signal is normalised to the background intensity for both cells in each image per timepoint. The signal intensity is shown as Z-intensity.

### Cell cycle length analysis

Cell cycle length was analysed for N2 and *weSi34; zapSi4*. Cell cycle length analysis was also done for N2 and sld-2*(zap13[sld-2EDDD])* for *wee1.3* RNAi. Time lapse z-stack movies were taken starting from pronuclei migration until the four-cell stage of the early embryo development in *C. elegans*. The time lapse interval was 8s. Cell cycle time of the AB cell was calculated as the time starting from pronuclei fusion until nuclear envelope breakdown of the AB cell. Movie was taken using a Deltavision microscope with Nomarski optics and a 60X oil objective. Cell cycle timing of the P1 cell was calculated as the time from pronuclei fusion until the nuclear envelope breakdown of the P1 cell.

### Progeny assays

N2, JA1563, PAZ3, *weSi34; zapSi3* and *weSi34; zapSi4* were synchronized by bleaching. Plates were kept at 25°C. L4 (P0) were singled on NGM plates seeded with OP50. After 24hrs P0 worm was transferred on new plate. This was done for five days until egg production stopped. The total production of fertilized eggs from each animal was calculated.

#### CRISPR *sld-3(2D)* and *sld-3(2E)* progeny analysis

Heterozygous hermaphrodites of sld-3(2D) and sld-3(2E) and homozygotes single site mutants were allowed to lay eggs at 20°C. Their larvae were singled on new plates and checked for sterility. After two days after reaching adulthood they were lysed and genotyped by PCR for the sld-3 locus.

### Embryonic lethality assays

The *sld-3* Crispr mutants (*sld-3(paz5), sld-3(paz6), sld-3(paz7), sld-3(paz8), sld-3(paz9), sld-3(paz10)*) were tested for embryonic lethality. N2 was used as a control. Plates were kept at 20°C. Young adults were singled on new plates and allowed to lay eggs for 24hrs. The hermaphrodite was then removed and the embryonic lethality was determined after additional 36 hours.

The *sld-2(8D); sld-3(2E)* MosSCI strain was tested for embryonic lethality. N2 and weSi34; zapSi4 were synchronized by bleaching. The plates were grown in 25°C until mid-J4 stage. Worms were singled on new plates and transferred again to new plates every 24hrs for five days until egg production stopped. Living larva and unhatched eggs were counted after additional 24hrs. Experiment was performed in 25°C.

## Acknowledgements

Thanks to Julie Ahringer for strains WM150 and KK571 and discussions. We thank Nathan Goehring for the worm strain KK1228 and strains KK300 and EU548 were obtained from the CGC. Work in the PZ lab was supported by AICR 10-0908, Wellcome Trust 107056/Z/15/Z and Gurdon Institute funding (Cancer Research UK C6946/A14492, Wellcome Trust 092096). MK was supported by HFSP grant LT000681/2017-L. AK was supported by Wellcome Trust studentship 203767/Z/16/Z.

## Supplementary Fig Legends

**Fig S1.** Analysis of single CDK site mutants in sld-3

**A.** Alignment of the CDK sites in Sld3/Treslin required for the interactions with Dpb11/TopBP1. The amino acid numbers of the two orthologous CDK sites in *C. elegans* SLD-3 are indicated below.

**B** and **C.** Box plots of progeny size (B) and embryonic lethality (C) of a wild type strain (N2) or strains with the CDK site T487 mutated to alanine, aspartic acid or glutamic acid by CRISPR.

**D** and **E.** As B/C except for the T438 CDK site.

**Fig S2.** Bypass of Sld3 and Sld2 phosphorylation is partially sufficient for cyclin E function.

**A.** Box plots of embryonic lethality from strains containing extra copies of *sld-2* / *sld-3*, both of which are inserted at a MosSCI sites and expressed from the *mex-5* promoter.

**B-E.** As A, but with RNAi of *cye-1* (B and C), *cyb-1* (D) or *cyb-3* (E).

**Fig S3.** SLD-3 and SLD-2 are asymmetrically localised in the early embryo

**A.** Box plot of SLD-3 mCherry signal ratio between the P1 and the AB cell.

**B.** Images from A. Images show brightfield, Hoescht and mCherry signal of the wild type (N2) or the respective MosSCI lines. mCherry signal corresponds to SLD-3 expression in the two-cell embryo. Scale bar is 10μm.

**C.** Quantitation of SLD-2-GFP fluorescence from live imaging. Graph shows fluorescent intensity signal of the AB and P1 cell. The graph represents one-minute time-lapse movies of two cell embryos. The time point 0 is the nuclear envelope breakdown (NEB) in the AB cell. Error bars are 95% CI; n = 8

**Fig S4.** Generation of PKC-3 interaction mutant in SLD-2.

**A.** Alignment of the C-termini of SLD-2 proteins from the indicated nematode species. The region of interaction between *C. elegans* SLD-2 and PKC-3 is indicated in red.

**B.** Summary of the yeast two-hybrid interactions between SLD-2 mutants and PKC-3 1-219.

**C.** Yeast two-hybrid growth assays of some of the mutants described in B.

D. Images of PKC-3 GFP and SLD-2 mCherry. This is the same image as Fig 6D, but with increased brightness of the mcherry signal to visualise SLD-2 in the cytoplasm. Scale bar is 10 μm.

